# Endothelin 2–Mediated Neuroinflammation Drives Hypertension-Associated Retinal Dysfunction

**DOI:** 10.64898/2026.01.26.701756

**Authors:** Yingkun Cui, Man-Ting Au, Lanlan Zhang, Ting Zhang, Li Pan, Hoi-Lam Li, Yuqi Zhang, Margaret MH Wu, Huan Li, Shing-Yan Roy Chung, Feng Pan, Chunyi Wen, Chi-Wai Do

**Author notes:** Contacts: Chi-wai Do; Address: HJ534, The Hong Kong Polytechnic University, Kowloon, Hong Kong, Chunyi Wen; Address:ST417, The Hong Kong Polytechnic University, Kowloon, Hong Kong.

## Abstract

Systemic hypertension is a significant risk factor for glaucoma, a leading cause of irreversible blindness, yet the mechanistic basis connecting the two conditions remains elusive. Here, we demonstrate that hypertensive rats develop accelerated retinal degeneration characterized by retinal thinning, functional impairment, reduced perfusion, and progressive neuroinflammation. Critically, these changes occur despite persistently lower intraocular pressure (IOP), indicating an IOP-independent mechanism of pathology. Transcriptomic and biochemical analyses identified a robust activation of the retinal endothelin system, with endothelin-2 (EDN2) being the earliest and most significantly upregulated component. AAV-delivered retinal *Edn2* knockdown completely restored retinal function, ameliorated neuroinflammation, and enhanced retinal blood flow in hypertensive rats. Pharmacological blockade of endothelin receptor type A (EDNRA), but not type B, replicated these protective effects without altering IOP. Our findings establish EDN2 as a central, IOP-independent driver of hypertension-associated retinal neurodegeneration and identify the endothelin-EDNRA axis as a potential therapeutic target to prevent vision loss in hypertensive patients.

## Introduction

Glaucoma, a leading cause of irreversible blindness worldwide (1) affecting over 76 million individuals (2), comprises a heterogeneous group of disorders characterized by progressive retinal ganglion cell loss and optic nerve degeneration (3). Although elevated intraocular pressure (IOP) is a major risk factor and current therapies primarily target IOP reduction (4–6), many patients continue to lose vision despite effective IOP control (7), implicating additional contributors to disease progression. Systemic hypertension (HT) is one such factor (8–11): epidemiological studies, including the Blue Mountains Eye Study, report a positive association between HT and primary open-angle glaucoma (POAG) (8), with higher blood pressure correlating with increased IOP and glaucoma prevalence (8–14). Nevertheless, the precise relationship among blood pressure, IOP, and glaucomatous neurodegeneration—and the underlying molecular mechanisms—remains incompletely defined.

Recent evidence suggests that the endothelin system, particularly endothelin-2 (EDN2), plays a key role as a mediator at the intersection of HT and glaucoma. Endothelin-1 (EDN1), a potent vasoconstrictor, is elevated in patients with essential HT and has been implicated in ocular vascular dysregulation and IOP modulation through endothelin receptors type A and B (EDNRA and EDNRB) (15–23). While the role of EDN1 in glaucoma has been extensively investigated, the contribution of endothelin-2 (EDN2)—another isoform with similar receptor affinity—remains poorly understood (24, 25). Notably, EDN2 expression is upregulated in the retina and optic nerve head in animal models of glaucoma (26). The contribution of EDN2 to retinal degeneration, particularly its impact on specific retinal cell populations such as alpha-retinal ganglion cells (αRGCs), which appear more susceptible to glaucoma damage, remains to be clarified (27, 28).

To address these gaps, we integrated human transcriptomic analyses with longitudinal studies in spontaneously hypertensive rats (SHR) and normotensive Wistar Kyoto rats (WKY) to investigate the effects of chronic HT on IOP, retinal structure and function, and neuroinflammation. We further examined the mechanistic role of EDN2 by employing targeted retinal knockdown (KD) and pharmacological blockade of endothelin receptors, assessing outcomes at molecular, cellular, and functional levels. This integrated approach enables the identification of early molecular events and cellular dysfunction in hypertension-induced retinopathy, emphasizing the interplay among BP, IOP, and EDN2 signaling. Collectively, these studies aim to delineate novel IOP-independent pathways of retinal degeneration and establish EDN2 as a potential therapeutic target for glaucoma and hypertensive retinal disease.

## Results

### Systemic hypertension accelerates IOP-independent retinal degeneration in rats

At the animal level, the impact of systemic hypertension on retinal integrity was investigated through longitudinal assessment of blood pressure, IOP, retinal function, and structure in SHR and age-matched WKY (Figures 1A-D). Systolic blood pressure (SBP) was significantly elevated in SHR throughout the study, exceeding WKY values by 26%, 47%, and 47%, at 4, 8, and 12 months, respectively (Figure 1E). Diastolic blood pressure (DBP) showed a parallel increase, reaching a 57% increase over WKY at 12 months (Figure 1F). Both SBP and DBP increased progressively with age in SHR, whereas levels remained relatively stable in WKY. Despite the elevated SHR, IOP was consistently but modestly lower in SHR than in WKY, with a mean difference < 1.4 mmHg across all time points (Figure 1G).

**Figure 1.**
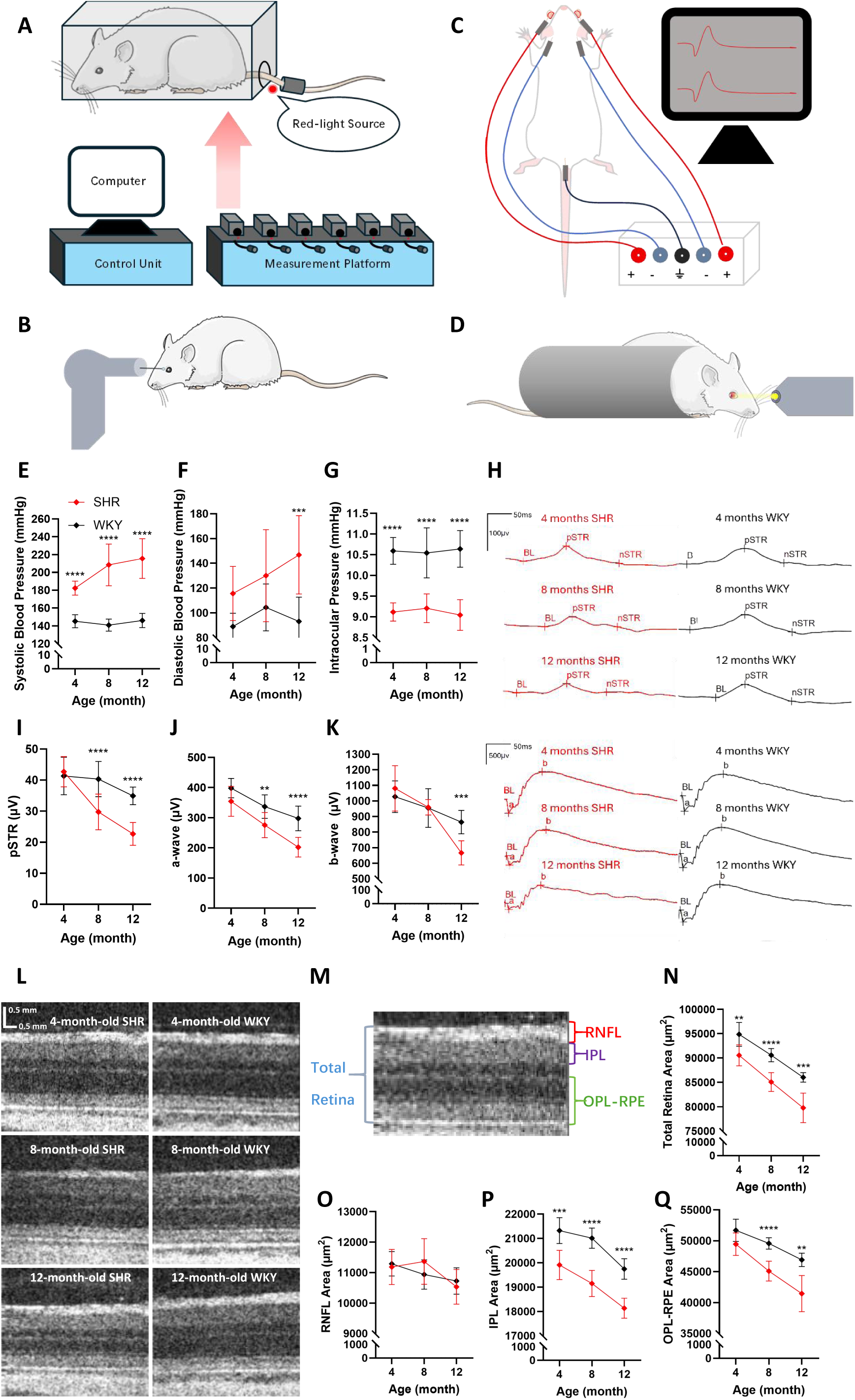
Systemic hypertension reduces functional retinal responses and retinal thickness. **A-D.** Illustration of the measurements of BP, IOP, ERG, and OCT, respectively; **E-G.** Measurements of SBP, DBP, and IOP in SHR and WKY (n=9 for SHR, n=9 for WKY); **H.** Representative ERG waveforms from 4, 8, and 12-month-old SHR and WKY; **I-K.** Measurements of pSTR amplitudes, a-wave, and b-wave amplitudes in SHR and WKY (n=9 for SHR, n=10 for WKY); **L.** Representative OCT images from 4, 8, and 12-month-old SHR and WKY; **M.** Identification of the RNFL, IPL, OPL-RPE, and total retina layers; **N-Q.** Measurements of total retinal, RNFL, IPL, and OPL-RPE areas in SHR and WKY (n=9 for SHR, n=9 for WKY). Data are shown as Mean ± SD. ** P < 0.01; *** P < 0.001; **** P < 0.0001 (Statistical significance was determined using a two-way ANOVA with Bonferroni’s multiple comparisons test for E-G, I-K, and N-Q).

Electroretinography (ERG) results revealed age-dependent declines in positive subthreshold response (pSTR), a-wave, and b-wave amplitudes in both groups, with more pronounced reductions in SHR (Figures 1H-K). Compared with WKY, SHR showed significant reductions in pSTR and a-wave amplitudes at 8 and 12 months and in b-wave amplitude at 12 months (Figures 1I-K). Quantitatively, SHR exhibited 27% and 19% reductions in pSTR and a-wave amplitudes at 8 months. By 12 months, these deficits increased to 40%, 29%, and 23% for pSTR, a-wave, and b-wave, respectively.

OCT revealed progressive retinal thinning in both strains (Figure 1L). Layer segmentation (RNFL, IPL, OPL–RPE, and total retina) is illustrated in Figure 1M. Total retinal thickness was consistently lower in SHR at all time points, decreasing by 4.6%, 6.1%, and 7.3% at 4, 8, and 12 months, respectively (Figure 1N). No significant difference in RNFL thickness was observed between the two groups (Figure 1O). In contrast, IPL thickness in SHR decreased by 6.6%, 8.9%, and 8.1% across the same intervals (Figure 1P), while the OPL-RPE complex was reduced by 4.3%, 9.0%, and 11.7% at corresponding time points (Figure 1Q).

### Systemic hypertension reduces retinal ganglion cell sensitivity and density with microglial activation

At the cellular level, light-evoked responses of αRGCs were recorded using the whole-cell patch-clamp electrophysiology in 8-month-old SHR and WKY to assess the effects of systemic hypertension on retinal function (Figure 2A). Both OFF- and ON-αRGCs were recorded (Figures 2B-C). Compared to WKY, αRGCs from SHR showed slower rising phases and delayed attainment of the full normalized response under equivalent light intensities (Figures 2D and F). Response thresholds were significantly elevated in SHR for both OFF- and ON-αRGCs, reflecting reduced light sensitivity. The average response threshold for OFF-αRGCs was 3.03 log units in SHR versus 2.12 log units in WKY (Figure 2E), while ON-αRGCs required 3.22 log units in SHR compared with 2.29 log units in WKY (Figure 2G).

**Figure 2.**
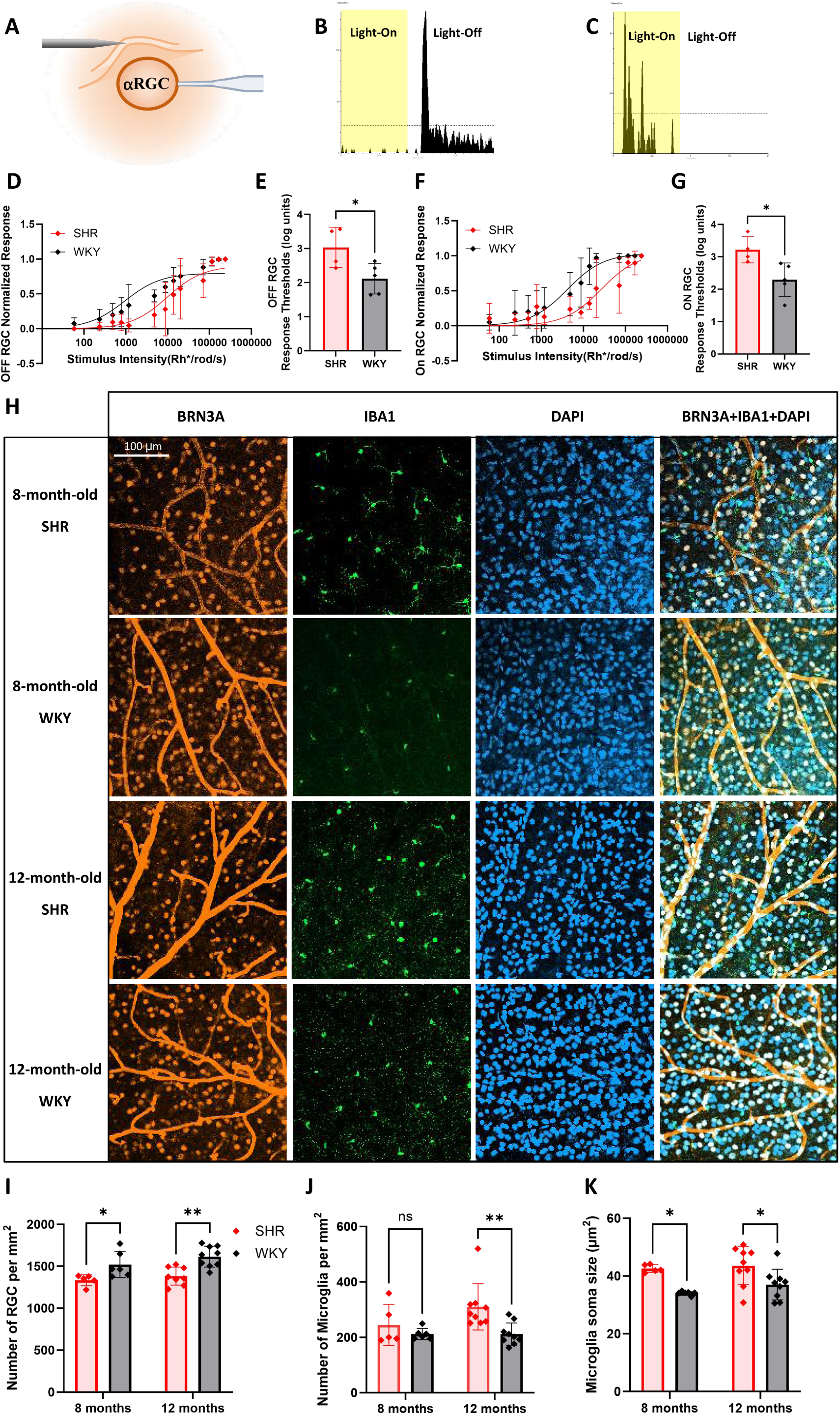
Systemic hypertension reduces RGC sensitivity and density, accompanied by microglial activation. **A.** Illustration of the patch clamp setup; **B.** Examples of characterized spikings from an OFF-αRGC; **C.** Examples of characterized spikings from an ON-αRGC; **D.** OFF-αRGC normalized responses at different stimulus intensities in 8-month-old SHR and WKY (n=4 for SHR, n=5 for WKY); **E.** OFF-αRGC response thresholds in SHR and WKY; **F.** ON-αRGC normalized responses at different stimulus intensities in SHR and WKY (n=4 for SHR, n=5 for WKY); **G.** ON-αRGC response thresholds in SHR and WKY; **H.** Representative images from the 8- and 12-month-old SHR and WKY whole-mounted retinas; **I.** Retinal ganglion cell counts (n=5 for 8-month-old SHR, n=6 for 8-month-old WKY, n=8 for 12-month-old SHR, n=9 for 12-month-old WKY); **J.** Microglial cell counts (n=5 for 8-month-old SHR, n=6 for 8-month-old WKY, n=9 for 12-month-old SHR, n=9 for 12-month-old WKY); **K.** Microglial soma size measurements (n=5 for 8-month-old SHR, n=6 for 8-month-old WKY, n=9 for 12-month-old SHR, n=9 for 12-month-old WKY). Data are presented as Mean ± SD. * P < 0.05; ** P < 0.01 (Statistical significance was determined using a two-way ANOVA with a two-tailed Student’s t-test for E and G, and a Bonferroni’s multiple comparisons test for I-K).

RGC and microglial populations were further analyzed using retinal whole-mounts from 8- and 12-month-old SHR and WKY (Figure 2H). RGCs were stained with an anti-BRN3A antibody, and microglia were stained with an anti-IBA1 antibody (29). RGC density was significantly lower in SHR than in WKY, decreasing by 12% at 8 months and 14% at 12 months (Figure 2I). Microglial density was comparable between groups at 8 months; but significantly elevated in SHR by 12 months (Figure 2J). Morphometric analysis revealed enlarged microglial soma in SHR, consistent with an activated phenotype. Soma area was 23% greater in SHR at 8 months and 16% greater at 12 months relative to WKY, both statistically significant (Figure 2K).

### Systemic hypertension reduces retinal blood supply

Retinal perfusion was assessed by quantifying the lumen areas of retinal arteries and veins. Across all ages, SHR showed markedly smaller arteries and venous lumens than WKY (Figure 3A). The measurement regions for arteries (red circles) and veins (blue circles) are shown in Figure 3B. The arterial lumen area in SHR decreased by 72%, 62%, and 77% relative to WKY at 4, 8, and 12 months, respectively, and decreased progressively with age in both groups (Figure 3C). At all time points, SHR showed smaller venous lumens, reduced by 70%, 69%, and 63% at 4, 8, and 12 months, respectively (Figure 3D).

**Figure 3.**
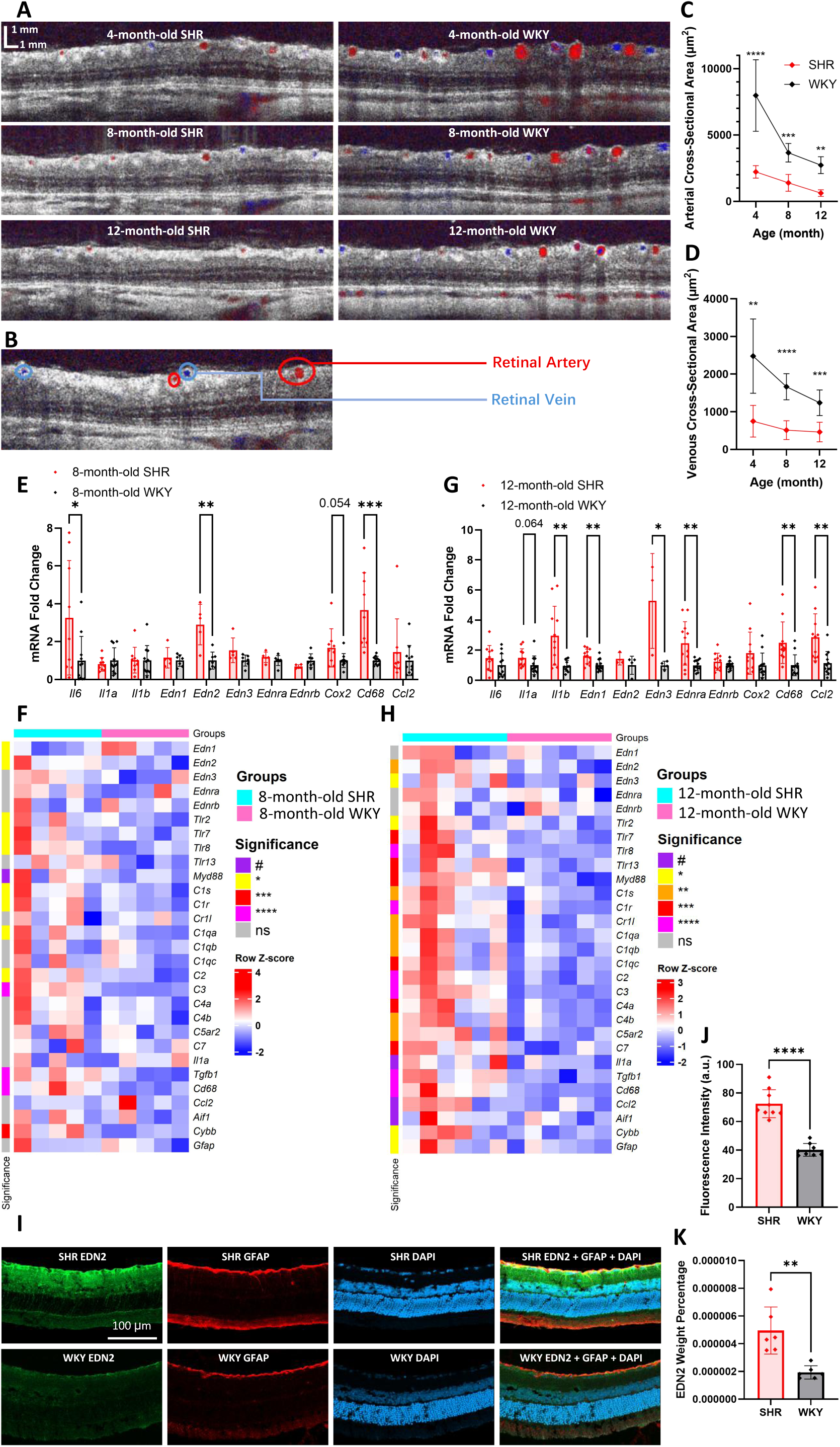
Systemic hypertension impairs retinal perfusion, induces neuroinflammation, and activates the retinal endothelin system. **A.** Representative images of arteries and venous lumen areas in 4-, 8-, and 12-month-old SHR and WKY; **B.** Schematic diagram indicating the measured arterial (red circle) and venous (blue circle) regions; **C-D.** Measurements of lumen areas of retinal arteries and veins in SHR and WKY (n=9/group); **E.** qPCR analysis of inflammatory markers at 8 months (n=5-9 for SHR, n=7-11 for WKY); **F.** Heatmap of selected target genes comparing 8-month-old SHR to WKY (n=5 for SHR, n=5 for WKY); **G**. qPCR results of retina samples from 12-month-old SHR and WKY (n=3-12 for SHR, n=4-13 for WKY); **H.** Heatmap of selected target genes comparing 12-month-old SHR to WKY (n=6 for SHR, n=6 for WKY); **I.** Representative images of retinal EDN2 and GFAP staining from 8-month-old SHR and WKY; **J.** Quantification of EDN2 fluorescence intensities (n=9 for SHR, n=8 for WKY); **K.** EDN2 weight percentage measured by ELISA (n=6 for SHR, n=6 for WKY). Data are shown as Mean ± SD. * P < 0.05; ** P < 0.01; *** P < 0.001; **** P < 0.0001 (Statistical significance was determined using a two-way ANOVA with Bonferroni’s multiple comparisons test for C-D, and a two-tailed Student’s t-test for others).

### Systemic hypertension activates the retinal endothelin system and triggers neuroinflammation

At the molecular level, quantitative PCR (qPCR) and RNA sequencing analyses were performed on retinas collected from 8- and 12-month-old SHR and WKY to assess the changes in the endothelin system and inflammatory gene expression profiles. The endothelin-related genes assessed included *Edn1*, *Edn2*, *Edn3*, Ednra, and Ednrb, while inflammation-associated targets encompassed cytokines (30–35), Toll-like receptors (TLRs) (36–38), and complement elements (26, 39) (Figure 3E-H). At 8 months, qPCR revealed elevated retinal expression of *Il6 a*nd *Cd68* in SHR compared with WKY. Within the endothelin system, only *Edn2* showed significant upregulation at this stage (Figure 3E). RNA sequencing data further demonstrated increased expression of complement components (*C2*, *C1r*, *C1s*, *C3*, and *C1qa*), TLRs (*Tlr2*, *Tlr7*, and *Tlr8*), and *Myd88* in SHR (Figure 3F). Inflammatory and microglial activation markers such as *Tgfb1*, *Cd68*, and *Cybb* were also significantly elevated. The Kyoto Encyclopedia of Genes and Genomes (KEGG) pathway analysis identified enrichment of neuroinflammation-associated pathways, including neutrophil extracellular trap formation, leukocyte transendothelial migration, and chemokine signaling, in 8-month-old SHR (Figure S1A). By 12 months, activation of both the endothelin system and inflammatory cascades became more pronounced. SHR showed significant upregulation of *Edn1*, *Edn2*, *Edn3*, and *Ednra* (Figure 3G and F). qPCR confirmed elevated expression of *Il1b*, *Edn1*, *Edn3*, *Cd68*, and *Ccl2* (Figure 3G). RNA sequencing revealed extensive elevation of neuroinflammatory and glaucoma-associated genes, including *Tgfb1*, multiple complement components (*C1qa*, *C1qb*, *C1qc*, *C1r*, *C1s*, *C2*, *C3*, *C4a*, *C4b*, *C5ar2*, *C7*), *Cd68*, TLRs (*Tlr2*, *Tlr7*, *Tlr8*, *Tlr13*), *Myd88*, *Cybb*, *Il1a*, *Gfap*, *Aif1,* and *Ccl2* (Figure 3H). Corresponding KEGG analyses highlighted significant enrichment of pathways linked to cytokine-cytokine receptor interactions, leukocyte transendothelial migration, complement and coagulation cascades, chemokine signaling, and TLR signaling (Figure S1B). Comparative pathway analyses revealed that complement and coagulation cascades were further upregulated in 12-month-old SHR relative to 8-month-old SHR, a change not observed in WKY (Figures S1C-D). Expression of *Edn2* and *Ednra* also increased with age in both strains (Figures S1E-F). Collectively, these findings indicate that systemic hypertension drives progressive activation of the retinal endothelin system and neuroinflammatory pathways, which intensify with age and are closely associated with molecular signatures of glaucoma pathogenesis.

### Systemic hypertension increases retinal expression of endothelin-2

SHR retinas showed markedly stronger EDN2 immunofluorescence compared with WKY (Figures 3I-J). Co-localization of EDN2 with GFAP, a key indicator of retinal astrocyte activation (40), revealed that retinal astrocytes are a principal source of EDN2 expression (Figure 3I). Quantitative ELISA confirmed significantly elevated EDN2 protein levels in SHR relative to WKY (Figure 3K), supporting hypertension-induced activation of the retinal endothelin system.

### Retinal endothelin-2 expression correlates with inflammation and aging in humans

To further investigate the association between *EDN2*, inflammation, and aging in humans, transcriptomic profiles from 322 retinal samples with mild age-related macular degeneration (AMD) (MG stages 1-2) were analyzed (GSE115828). *EDN2* expression showed a modest but significant positive correlation with donor age (P = 0.0081, r = 0.147, Figures S2A-B). Notably, *EDN2* levels correlated positively with several complement components (*C1R, C1QA, C1QB, C1QC, C4A, C4B*) and inflammatory markers, including *CD68, GFAP, TGFB1*, and *IL6* (Figure S2C). These findings corroborate the experimental data in SHR retinas, suggesting that systemic hypertension promotes retinal neuroinflammation through EDN2 upregulation, potentially triggering complement activation and inflammatory cascades.

### Edn2 knockdown improves retinal function and perfusion in hypertensive rats

The experimental protocol for AAV-mediated *Edn2* KD and subsequent assessments is illustrated in Figure 4A. The efficacy of *Edn2* silencing at the protein level was confirmed by ELISA and IHC analyses. EDN2 immunofluorescence intensity was markedly reduced in SHR treated with *Edn2* KD AAV compared with those receiving control virus (Figures 4B-C). In WKY, *Edn2* KD AAV produced a nonsignificant trend toward lower EDN2 expression (Figures 4B and D). ELISA quantification corroborated these findings, revealing significant reductions in retinal EDN2 protein levels following *Edn2* KD AAV treatment in both SHR and WKY (Figures 4E-F).

**Figure 4.**
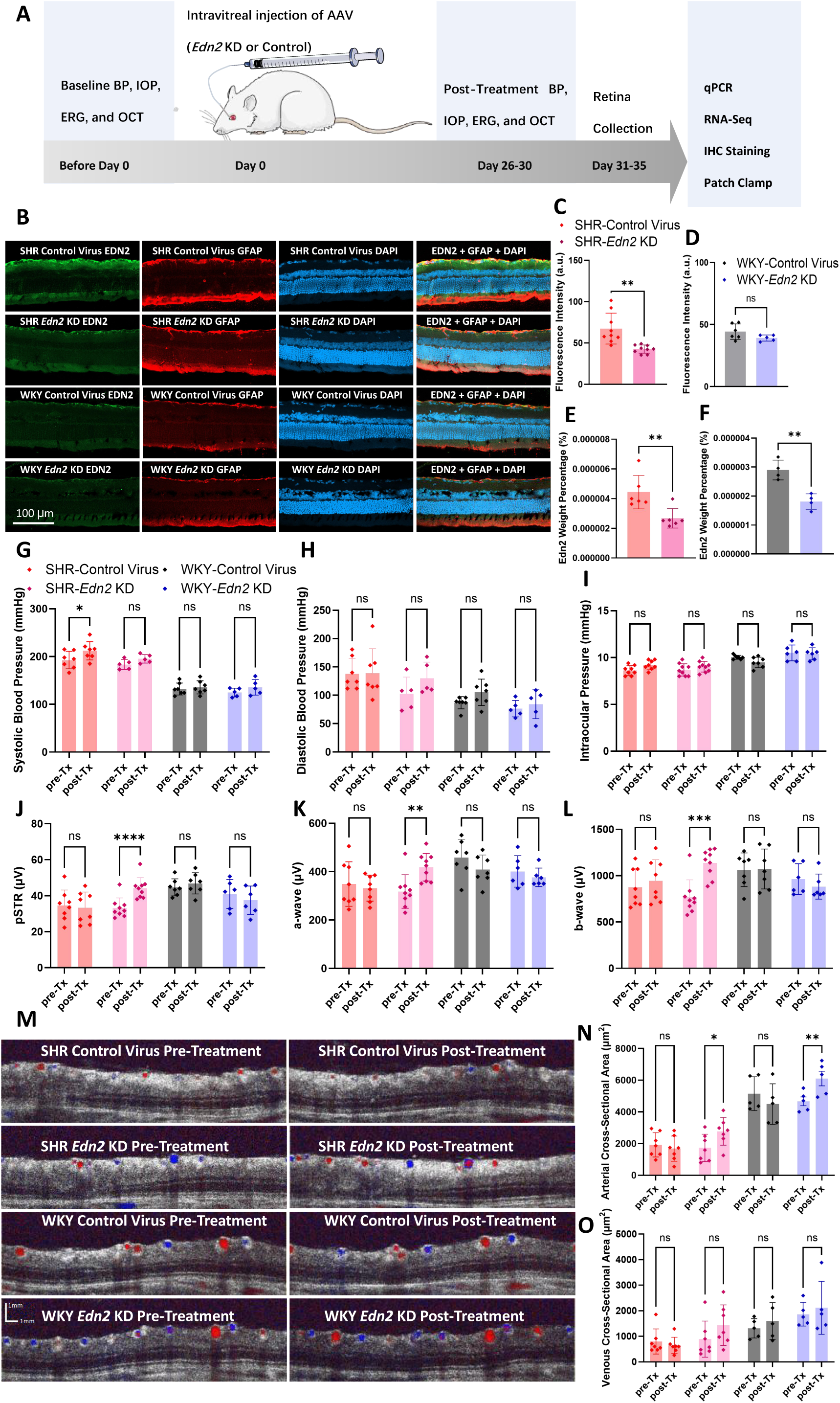
Retinal *Edn2* KD rescues functional responses and restores perfusion in SHR. **A.** Experimental protocol for AAV treatment and measurements; **B.** Representative images of retinal cryosections from SHR and WKY injected with *Edn2* KD AAV or control AAV; **C-D**. Quantification of EDN2 fluorescence intensities in SHR and WKY (n=9 for SHR-Control Virus, n=9 for SHR-*Edn2* KD, n=6 for WKY-Control Virus, n=5 for WKY-*Edn2* KD); **E-F**. EDN2 protein levels measured by ELISA (expressed as a percentage of total protein) in SHR and WKY (n=6 for SHR-Control Virus, n=6 for SHR-*Edn2* KD, n=4 for WKY-Control Virus, n=4 for WKY-*Edn2* KD); **G-H.** Pre- and post-treatment measurements of SBP and DBP in SHR and WKY (n=7 for SHR-Control Virus, n=5 for SHR-*Edn2* KD, n=7 for WKY-Control Virus, n=5 for WKY-*Edn2* KD); **I-L.** Pre- and post-treatment measurements of IOP, pSTR, a-wave, and b-wave in SHR and WKY (n=8 for SHR-Control Virus, n=9 for SHR-*Edn2* KD, n=7 for WKY-Control Virus, n=6 for WKY-*Edn2* KD); **M.** Representative images of pre- and post-treatment lumen areas of retinal arteries and veins from SHR and WKY; **N-O.** Pre- and post-treatment measurements of lumen areas of retinal arteries and veins in SHR and WKY (n=7 for SHR-Control Virus, n=7 for SHR-*Edn2* KD, n=5 for WKY-Control Virus, n=5 for WKY-*Edn2* KD). Pre-Tx refers to pre-treatment, and Post-Tx refers to post-treatment. Data are shown as mean ± SD. * P < 0.05; ** P < 0.01; *** P < 0.001; **** P < 0.0001 (Statistical significance was determined using a two-tailed Student’s t-test for C-F, and a two-way ANOVA with Bonferroni’s multiple comparisons test for G-L and N-O).

Physiological assessments indicated that *Edn2* KD AAV had no significant effect on SBP, DBP, and IOP in SHR (Figure 4G-I), although control AAV induced a mild SBP increase in SHR (Figure 4G). Neither AAV treatment influenced these parameters in WKY. Importantly, *Edn2* KD significantly improved retinal function in SHR, with treated eyes showing 33%, 29%, and 48% increases in pSTR, a-wave, and b-wave amplitudes, respectively, compared with control AAV (Figures 4J-L). Control AAV had no significant effect on retinal responses in SHR, and *Edn2* KD did not affect retinal function in WKY. Retinal thickness remained unchanged across all conditions (data not shown).

Vascular analysis revealed enhanced retinal perfusion following *Edn2* KD. The arterial lumen area increased by 61% in SHR and 31% in WKY relative to control AAV-treated eyes (Figures 4M-N). Although the venous lumen area showed a 62% increase in SHR after *Edn2* KD, this difference approached but did not reach statistical significance (P = 0.1580). No significant vascular changes were observed in WKY (Figure 4O). Taken together, these results demonstrate that suppression of *Edn2* expression restores retinal function and improves vessel caliber in hypertensive rats without affecting systemic or IOP.

### Edn2 knockdown restores retinal ganglion cell sensitivity and density and reduces microglial activation in hypertensive rats

*Edn2* KD AAV treatment accelerated the response kinetics and improved light sensitivity in SHR αRGCs. OFF-αRGCs from *Edn2* KD-treated SHR reached full normalized responses more rapidly and showed significantly lower response thresholds (2.91 log units) compared with control-AAV-treated SHR (3.29 log units) (Figures 5A-B). Similarly, ON-αRGCs in SHR showed reduced thresholds following *Edn2* KD (1.89 log units) relative to control treatment (2.87 log units) (Figures 5D-E). Conversely, WKY αRGCs treated with *Edn2* KD AAV displayed slower response kinetics and nonsignificant increases in response thresholds for both OFF- and ON-αRGCs compared to control AAV (Figures 5A, C-D, and F). Specifically, thresholds were 2.69 log units versus 2.29 log units for OFF-αRGCs (Figure 5C) and 2.54 log units versus 1.84 log units for ON-αRGCs (Figure 5F), differences that did not reach statistical significance.

**Figure 5.**
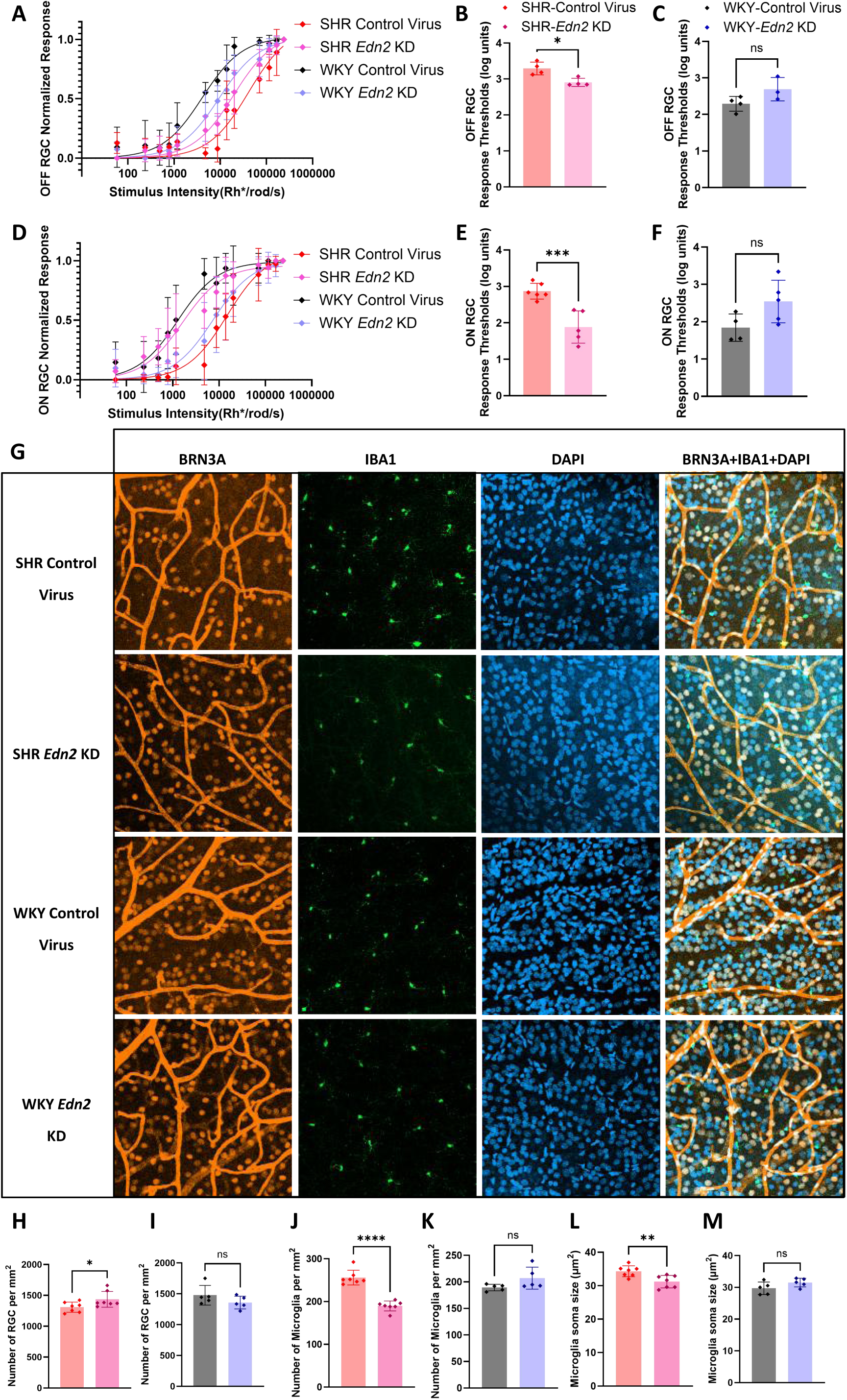
Retinal *Edn2* KD restores RGC sensitivity and density, with reduced microglial activation, in SHR. **A.** Post-treatment OFF-αRGC normalized responses at different stimulus intensities in SHR and WKY (n=4 for SHR-Control Virus, n=4 for SHR-*Edn2* KD, n=4 for WKY-Control Virus, n=3 for WKY-*Edn2* KD); **B-C.** Post-treatment OFF-αRGC response thresholds in SHR and WKY; **D.** Post-treatment ON-αRGC normalized responses at different stimulus intensities in SHR and WKY (n=6 for SHR-Control Virus, n=5 for SHR-*Edn2* KD, n=4 for WKY-Control Virus, n=5 for WKY-*Edn2* KD); **E-F.** Post-treatment ON-αRGC response thresholds in SHR and WKY; **G.** Representative images from post-treatment SHR and WKY retinas; **H-I.** Retinal ganglion cell counts from post-treatment SHR and WKY retinas; **J-K.** Microglial cell counts from post-treatment SHR and WKY retinas; **L-M.** Microglial soma size measurements from post-treatment SHR and WKY retinas. Data are shown as mean ± SD. * P < 0.5; ** P < 0.01; **\*\*\*** P < 0.001; **** P < 0.0001 (Statistical significance was determined using a two-tailed Student’s t-test for B-C, E-F, and H-M)

*Edn2* KD AAV significantly increased RGC density in SHR retinas compared with control AAV treatment (Figures 5G-H). Moreover, *Edn2* KD reduced both microglial cell density and soma size in SHR (Figures 5J and L), consistent with suppression of retinal microglial activation. No significant differences in RGC or microglial parameters were observed between *Edn2* KD and control-treated WKY retinas (Figures 5I, K, and M). These findings demonstrate that downregulation of *Edn2* restores retinal ganglion cell responsiveness and survival while dampening microglial activation in hypertensive retinas.

### Edn2 knockdown attenuates retinal neuroinflammation in hypertensive rats

In SHR, AAV-mediated *Edn2* KD not only reduced *Edn2* expression but also downregulated multiple endothelin and inflammatory genes, including *Edn3*, *Ednrb*, *Il1a*, *Il1b*, *C3*, *C1qa*, and *Tnfa* (Figure 6A). KEGG pathway analysis revealed significant suppression of complement and coagulation cascades as well as cytokine-cytokine receptor interaction pathways (Figure 6B). RNA sequencing confirmed broad downregulation of neuroinflammatory genes, including complement components (*C1r*, *C1s*, *C1qa*, *C1qb*, *C1qc*, *C2*, *C3*, *C4a*, *C4b*, *C5ar2*), TLRs (*Tlr2*, *Tlr7*, *Tlr8*, *Tlr13*), and microglial or inflammatory activation markers (*Aif1*, *Cd68*, *Gfap*, *Tgfb*1, *Myd88*, *Ccl2*, *Cybb*) (Figures 6C-D). In WKY, *Edn2* KD AAV significantly decreased *Edn2* mRNA levels (Figures 6E, G-H), though *Edn1* expression modestly increased (Figure 6E). KEGG pathways analysis supported suppression of complement and cytokine signaling pathways (Figure 6F). Transcriptomic profiling identified downregulation of *Edn3*, *Ednrb*, complement components (*C1qa*, *C1qc*, *C1r*, *C3*, *C1s*, *C4a*, *C4b*), and inflammatory markers (*Gfap*, *Tlr7*, *Tlr8*, *Aif1*, *Cyb*b, *Tnfa*) (Figure 6H). Importantly, the extent of transcriptomic alteration was substantially smaller in WKY than in SHR (Figure 6G), suggesting that *Edn2* KD preferentially mitigates hypertension-associated retinal inflammation.

**Figure 6.**
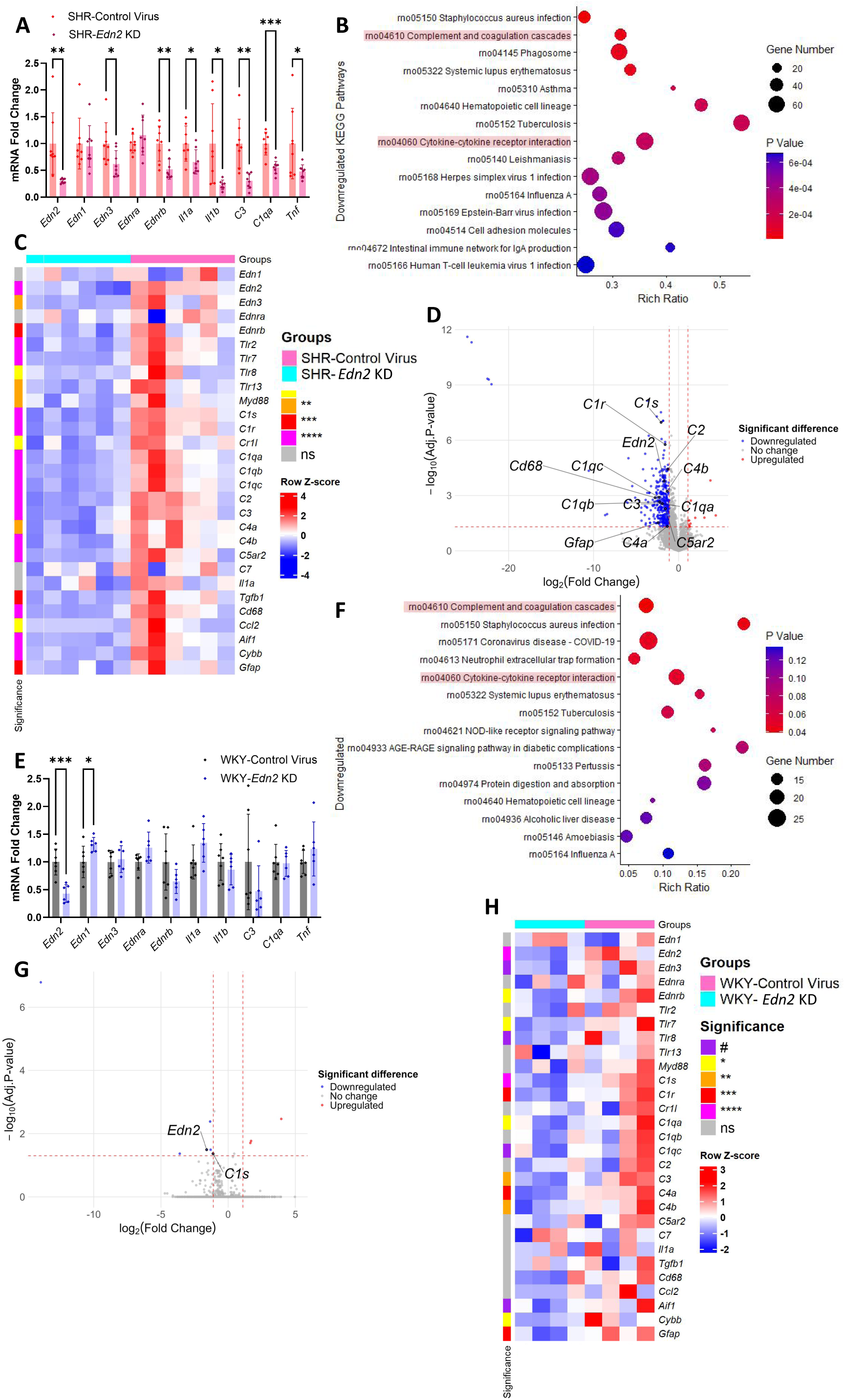
Retinal *Edn2* KD suppresses neuroinflammation in SHR. **A.** qPCR results of SHR injected with either *Edn2* KD AAV or control AAV (n=8 for SHR-Control Virus, n=7-8 for SHR-*Edn2* KD); **B.** Downregulated KEGG pathways in SHR injected with *Edn2* KD AAV compared to those injected with control AAV (n=6 for SHR-Control Virus, n=6 for SHR-*Edn2* KD); **C.** Heatmap of selected target genes comparing SHR receiving *Edn2* KD virus against SHR receiving control virus; **D.** Volcano plots showing downregulated targets in SHR treated with *Edn2* KD AAV; **E.** qPCR results of WKY injected with either Edn2 KD AAV or control AAV (n=7 for WKY-Control Virus, n=6 for WKY-*Edn2* KD); **F.** Downregulated KEGG pathways in WKY injected with *Edn2* KD AAV compared to those injected with control AAV (n=4 for WKY-Control Virus, n=4 for WKY-*Edn2* KD); **G.** Volcano plots illustrating downregulated targets in WKY treated with *Edn2* KD AAV; **H.** Heatmap of selected target genes comparing WKY receiving *Edn2* KD virus against WKY receiving control virus. Data are presented as Mean ± SD. * P < 0.5; ** P < 0.01; *** P < 0.001 (Statistical significance was determined using a two-tailed Student’s t-test for A and E).

### Pharmacological blockade of the endothelin type A receptor, but not the type B receptor, restores retinal function in hypertensive rats

To further investigate the downstream receptor responsible for EDN2-mediated retinal dysfunction, selective blockade of the EDNRA and EDNRB was performed in SHR and WKY via intravitreal injection of the antagonists BQ123 and BQ788, respectively. The effects of these treatments on IOP and retinal function are summarized in Figure 7. Neither BQ123 nor BQ788 altered IOP in SHR. In WKY, BQ123 modestly reduced IOP in the treated eye, from a baseline of 10.0 mmHg to 9.3 mmHg at Day 8 and 9.1 mmHg at Day 18 (Figure 7B). Importantly, BQ123 treatment significantly improved retinal function in SHR but not in WKY. In SHR, the pSTR amplitude increased by 32% and 58% relative to baseline on Day 9 and Day 19 post-injection, respectively (Figure 7C). The a-wave and b-wave amplitudes were also enhanced, increasing by 20% and 21% at Day 19 (Figures 7D-E), No significant functional changes were observed in WKY after BQ123 treatment. Conversely, BQ788 did not produce significant changes in retinal responses in either strain (Figures 7C-E).

**Figure 7.**
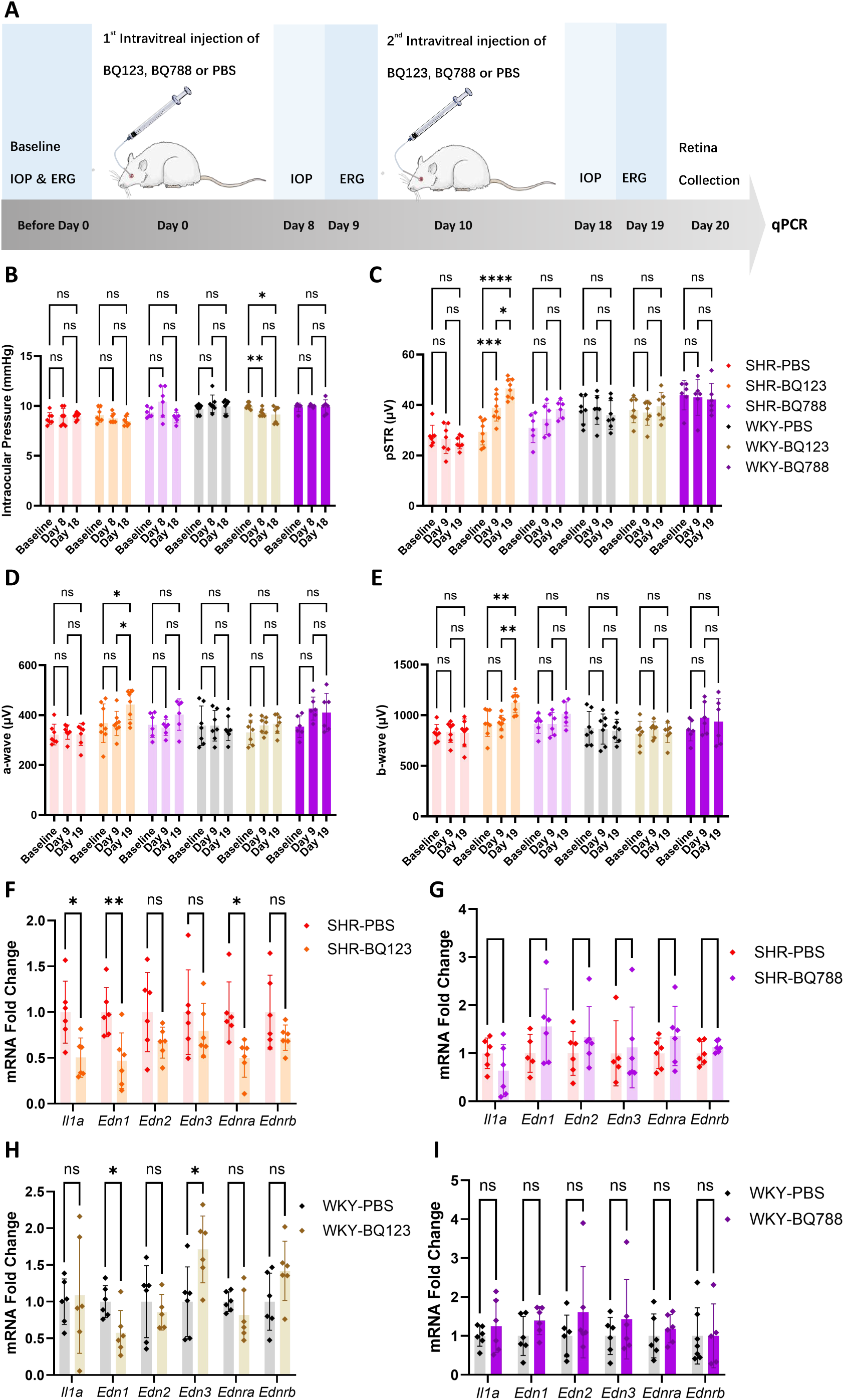
Selective blockade of EDNRA, but not EDNRB, rescues retinal function in SHR. **A.** Experimental design for treatment with BQ123 and BQ788 in SHR and WKY; **B-E.** IOP and functional retinal responses of SHR and WKY following injections of BQ123, BQ788, or PBS (n=7 for SHR-PBS, n=7-8 for SHR-BQ123, n=6 for SHR-BQ788, n=7 for WKY-PBS, n=7 for WKY-BQ123, n=6 for WKY-BQ788); **F-G.** qPCR results for SHR injected with BQ123 or BQ788 compared to SHR injected with PBS (n=5-6 for SHR-PBS, n=6 for SHR-BQ123, n=6 for SHR-BQ788); **H-I.** qPCR results for WKY injected with BQ123 or BQ788 compared to WKY injected with PBS (n=6-7 for WKY-PBS, n=5-6 for WKY-BQ123, n=5-6 for WKY-BQ788). Data are presented as Mean ± SD. * P < 0.05; ** P < 0.01; *** P < 0.001; **** P < 0.0001 (Statistical significance was determined using a two-way ANOVA with Bonferroni’s multiple comparisons test for B-E, while for F-G the statistical significance was determined using a two-tailed Student’s t-test).

At the molecular level, SHR retinas treated with BQ123 showed reduced expression of *Il1a*, *Edn1*, and *Ednra* (Figure 7F). In WKY, BQ123 significantly downregulated *Edn1* but induced a mild upregulation of *Edn3* (Figure 7H). BQ788 treatment did not significantly alter gene expression in either SHR or WKY (Figures 7G and I).

## Discussion

In this study, we demonstrated that SHR develop significant retinal dysfunction and RGCs’ loss despite exhibiting lower IOP compared to WKY controls. By integrating functional, structural, and molecular assessments, we found that impaired retinal blood supply and heightened neuroinflammation, rather than elevated IOP, are the principal drivers of retinal degeneration in an inherited hypertension rat model.

Noteworthily, we identified EDN2 as a central mediator of these pathological processes. EDN2 is upregulated early in hypertension-induced retinal injury and orchestrates both vascular dysregulation and neuroinflammatory responses. Genetic ablation of *Edn2* or pharmacological blockade of its receptor, EDNRA, robustly restored retinal function and structure in SHR, without influencing BP or IOP, highlighting the endothelin-EDNRA axis as a promising therapeutic target.

Our study advances current understanding of pathomechanism linking hypertension and retinal degeneration via a detailed characterization in our unique SHR model. Consistent with previous reports, SHR exhibited elevated BP but paradoxically showed statistically significantly lower IOP compared to WKY controls. This reduction in IOP was not attributable to differences in central corneal thickness (Figures S3A-B), suggesting alternative mechanisms such as vascular rarefaction and hyalinization in the ciliary body (41). Importantly, the mild difference in IOP (approximately 1 mmHg) between groups should not be considered clinically significant. Therefore, retinal degeneration in SHR was induced by biochemical changes rather than by mechanical compression.

To further elucidate the impact of blood pressure on retinal function, we performed correlation analyses between SBP/DBP and ERG amplitudes. Our results revealed SBP as a more sensitive predictor of retinal dysfunction than DBP, implicating elevated SBP as a key driver of retinal injury in HT. Both SBP and DBP showed significant negative correlations with retinal functions at 12 months, but these were not observed at a young age in the developmental and maturation stages. It might suggest that hypertension exacerbates functional decline with systemic aging and degeneration (Figures S4-S5). This observation was consistent with a previous human study, which indicated that systemic hypertension worsened glaucoma in the elderly population but not in young patients (42). At the cellular level, we provide the first evidence that chronic HT impairs light-evoked responses in both ON- and OFF-αRGCs, with *Edn2* KD selectively improving ON-αRGC sensitivity. This finding aligns with previous reports of ON-αRGC vulnerability in other retinal diseases (43) and suggests subtype-specific plasticity in response to injury.

Mechanistically, transcriptomic analyses revealed that EDN2 promotes retinal neuroinflammation by activating complement components, a process corroborated by positive correlations in human retinal transcriptomic datasets. *Edn2* KD markedly downregulated complement components and inflammatory markers while increasing retinal blood flow in SHR, with less pronounced effects in WKY. The differential response to *Edn2* KD between SHR and WKY suggests that EDN2 serves as a critical mediator through which hypertension exacerbates retinal neurodegeneration.

Complement activation is recognized as an early event in glaucoma and other neurodegenerative diseases, and its inhibition is protective in animal models (26, 39, 44). Our data support a model in which EDN2 exacerbates HT-induced retinal degeneration through complement-mediated neuroinflammatory pathways, independent of IOP elevation. Moreover, pharmacological inhibition of EDNRA (BQ123), but not EDNRB (BQ788), restored retinal function in SHR, including pSTR recovery and improvements in a- and b-wave amplitudes, without influencing IOP. No functional improvements were observed in normotensive WKY. These strain-specific effects highlight EDN2 signaling through EDNRA as a key IOP-independent driver of HT-induced retinal neurodegeneration.

Our study has several strengths, including the use of longitudinal in vivo assessments, integration of functional, structural, and molecular analyses, and the application of both genetic and pharmacological interventions targeting EDN2 signaling over a prolonged period. Nonetheless, our study has limitations. First, the current analyses focus on global retinal improvements, but the precise mechanistic cellular interactions among different retinal cell types through which EDN2 promotes RGC survival remain to be elucidated. Further studies are warranted to delineate the cell-type–specific contributions of EDN2 and to investigate its role in other models of retinal neurodegeneration. Second, our work focused on the influence of hypertension on the retina at stages when visual function is already affected. How hypertension impacts ocular growth and development during earlier stages has not yet been investigated. A comprehensive atlas documenting the impact of hypertension on the eye across the lifetime is required and deserves further study.

In conclusion, we identify EDN2 as a central mediator of HT-induced retinal degeneration, acting through IOP-independent hemodynamic disruption and neuroinflammatory mechanisms. Both genetic and pharmacological inhibition of EDN2 signaling restores retinal function and structure in SHR, highlighting EDN2 as a promising therapeutic target for hypertensive retinopathy and potentially for glaucoma. These findings underscore the importance of targeting early molecular events in disease progression and provide a framework for future studies aimed at developing novel neuroprotective interventions against retinal neurodegeneration.

## Methods

### Animals

SHR and WKY were used as hypertensive and normotensive controls, respectively. All animals were housed in a temperature-controlled environment (23 ± 1°C) under a 12-hour light/dark cycle, with free access to food and water. The BP, IOP, retinal function, structure, and hemodynamic parameters were assessed longitudinally at 4, 8, and 12 months of age in both groups. Sex was not considered a biological variable in this study. Both male and female animals were included because similar ocular phenotypes were observed in both sexes.

### Measurement of Blood Pressure and Intraocular Pressure

The SBP and DBP in SHR and WKY were measured using the BP-2000 Blood Pressure Analysis System (Visitech Systems, USA) (Figure 1A). Measurements were performed between 2 pm and 5 pm to minimize circadian variability. The IOP was measured using a rebound tonometer (TonoLab, Icare, Finland) between 1 pm and 3 pm (Figure 1B). Six readings were averaged for each eye. The measurement sequence was randomized between SHR and WKY groups, and all recordings were obtained in awake animals.

### Measurements of Functional Retinal Responses

Functional retinal responses in SHR and WKY were recorded using a full-field ERG (Ganzfeld, Q450; RETI Animal, Roland Consult, Germany) (Figure 1C). The animals were dark-adapted for at least 6 hours before data acquisition. Prior to recordings, rats were anesthetized with a ketamine-xylazine mixture, and pupils were dilated with 0.5% tropicamide (Mydrin-P, Santen Pharmaceutical, Japan). RGC activity was assessed via the positive scotopic threshold response (pSTR) (45), recorded at a stimulus intensity of -4.8 log cd·s/m². Photoreceptors and inner retinal activities were represented by the a-wave (46) and b-wave (47), respectively. The amplitudes of the a- and b-waves were measured under single-flash stimulation at 1.3 log cd·s/m². All recordings were acquired with a bandpass filter of 0.1–1000 Hz.

### Measurements of Retinal Thickness and Hemodynamics using OCT

The retinal morphology was imaged using an Optical Coherence Tomography (OCT) system (Envisu-R2210, Bioptigen, Leica Microsystems, IL, USA) (Figure 1D). Retinal thickness was measured radially from the center of the optic nerve at a radius of 1 mm. Parameters included the total retina retinal thickness as well as the thicknesses of the RNFL, IPL, and OPL-RPE complex. Each OCT scan consisted of 100 images, of which eight (1^st^, 13^th^, 26^th^, 38^th^, 51^st^, 63^rd^, 76^th^, and 88^th^) were selected for analysis. When the selected frames overlapped with major retinal vessels, adjacent images were used to minimize vascular interference in RNFL measurements. Thickness analysis was performed manually with ImageJ. To avoid the topographic influence of the optic nerve head, all measurements excluded the central 0.5 mm radius around its center. Mean values from the eight selected images were used to determine the final thicknesses.

The lumen area of retinal arteries and veins was quantified using the Doppler mode of the same OCT system (48, 49). Circle scans centered on the optic nerve head with a 0.25 mm radius were performed, each acquired with 50 repeated scans. The retinal arteries and veins supplying the inner retina near the optic disc were identified (Figure 3B); red signals indicated arterial and blue signals indicated venous blood flow (45). Analysis was performed manually using ImageJ. For each Doppler scan, 50 frames were captured, and 10 evenly spaced images (6^th^, 11^th^, 16^th^, 21^st^, 26^th^, 31^st^, 36^th^, 41^st^, and 46^th^) were selected for analysis. The average values from these 10 frames were used to calculate the final arterial and venous lumen areas.

### Measurements of light-evoked spike activity of αRGCs

Retinas from 8-month-old SHR and WKY were dissected and mounted in a patch-clamp recording chamber for extracellular measurement of light-evoked spike activity in αRGCs (Figure 2A). These αRGCs were identified based on their large somata in the ganglion cell layer, characteristic light responses, and subsequent Neurobiotin filling, which confirmed their distinctive morphology. Post-treatment retinas from AAV-injected eyes were analyzed similarly. During recordings, retinas were stimulated with green light (525 nm) at intensities ranging from 29 to 8.4 × 10⁵ Rh∗ rod⁻¹ s⁻¹. Spike activity from both OFF- and ON-type αRGCs was captured, and the representative spiking patterns from OFF-αRGC and an ON-αRGC are shown in Figures 2B and 2C. Normalized responses were plotted against light intensity and fitted to the Michaelis-Menten equation (Figures 2D, 2F, 5A, and 5D). Light sensitivity threshold was defined as the stimulus intensity required to elicit 5% of the maximum response amplitude.

### Tissue processing and Immunohistochemical staining

For immunostaining of RGCs and microglia, eyes were fixed in 4% PFA for 2 hours at room temperature, and retinas were carefully isolated for whole-mount staining. RGCs were labeled with an anti-BRN3A antibody (1:400; MAB1585, Sigma Millipore), and microglia were labeled with an anti-IBA1 antibody (1:800; 019-19741, Wako). Following primary antibody incubation, retinas were counterstained with DAPI and flat-mounted under coverslips. Images were acquired using an upright confocal microscope (LSM 800, Zeiss) with a 20x magnification. For each retina, 24 fields were imaged. Microglial number and soma size were quantified manually using ImageJ, and RGCs were counted manually. For cryosection immunostaining, eyes were embedded and cryosectioned at a thickness of 7 µm, and sections within 1 mm of the optic nerve head were collected. Sections were labeled with anti- EDN2 (1:350; MBS9234016, Mybiosource) and anti-GFAP (1:350; 14-9892-82, eBioscience) antibodies. Five representative images were obtained per slide using the same confocal imaging system.

### The RNA Sequencing and qPCR

Total RNA was extracted from retinas using the Quick-RNA Miniprep Kit (Zymo Research, USA) and stored at -80 °C until analysis. RNA sequencing was performed by the Beijing Genomics Institute (BGI). Samples were sequenced using the DNBSEQ platform with a paired-end read length of 100 bp (PE100). Reads of low quality, with adaptor sequences, or high levels of N base were filtered using SOAPnuke (v1.5.6) (50). Bowtie2 (v2.4.4) was used to align the clean reads to the reference genes in the genome (GCF_000001895.5_Rnor_6.0) (51). Gene expression was quantified using RSEM (v1.2.28) (52). Differential expression analysis of gene expression was performed using DESeq2 (v1.40.2) with Benjamini-Hochberg correction (53). KEGG enrichment analysis was performed on Differential Expression Genes (DEGs) with adjusted p-value less than 0.1 using GAGE (v2.50.0) (54). Four comparisons were analyzed: 8-month-old SHR and WKY; 12-month-old SHR and WKY; SHR injected with *Edn2* KD AAV and SHR injected with control AAV; and WKY injected with *Edn2* KD AAV and WKY injected with control AAV. The top 15 upregulated pathways were shown for SHR and WKY comparisons, and the top 15 downregulated pathways for AAV comparisons. Genes in cytokine–cytokine interaction and complement and coagulation cascades were studied, with selected differentially expressed genes of interest plotted using ggplot2 (v4.0.1) and ComplexHeatmap (v2.16.0) in RStudio.

For qPCR, cDNA was synthesized from 1 μg total RNA using the High-Capacity cDNA Reverse Transcription Kit (Thermo Fisher Scientific, USA). The mRNA levels of inflammation- and endothelin-related targets including *Il6*, *Tnfa*, *Il1a*, *Il1b*, *Ccl2*, *Cox2*, *Cd68*, *C1qa*, *C3*, *Edn1*, *Edn2*, *Edn*3, *Ednra*, and *Ednrb*) were quantified, with acidic ribosomal phosphoprotein P0 (*36b4*) as the housekeeping gene. Primer sequences are listed in Table 1.

**Table 1.**
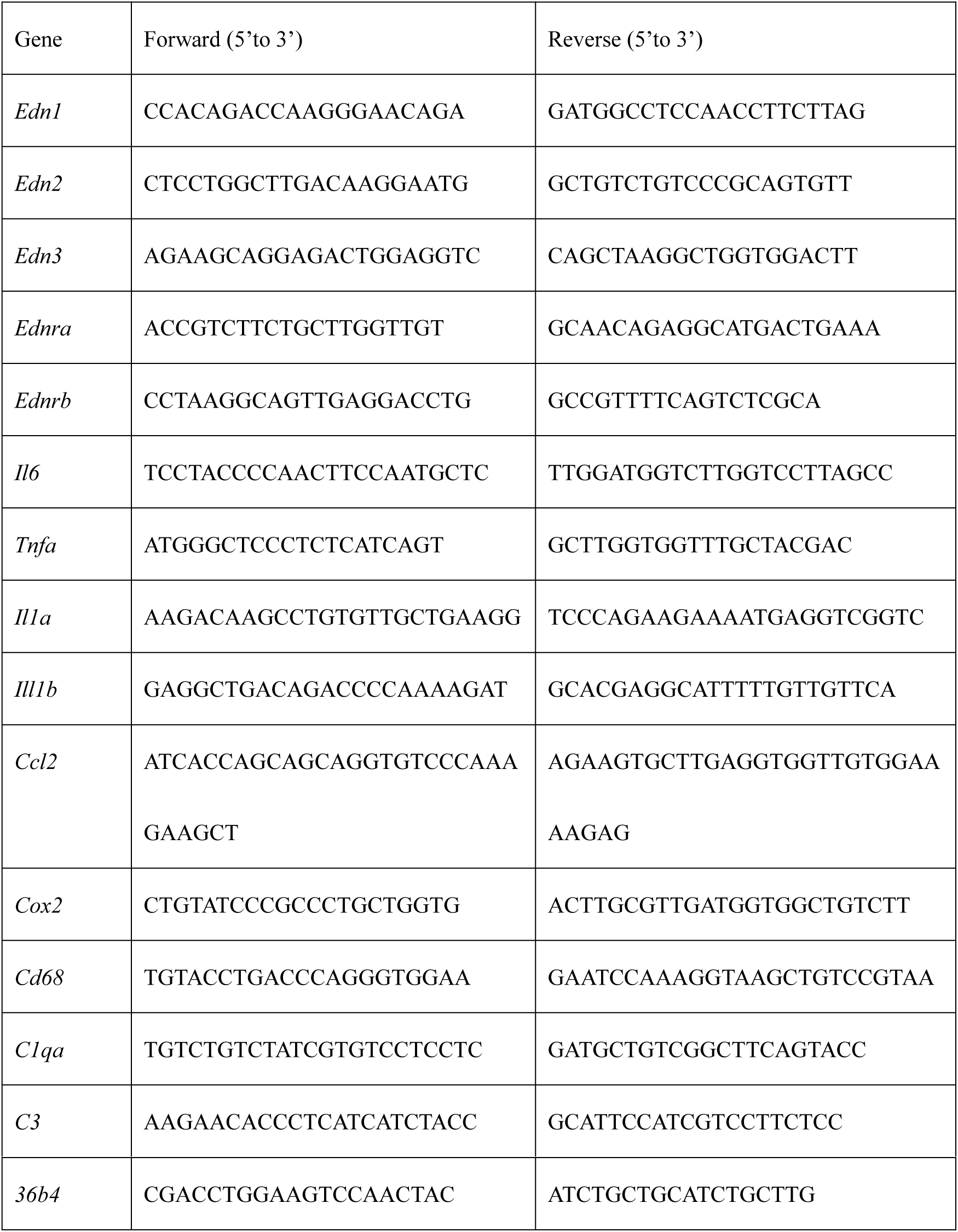
The sequences of the qPCR primers.

### Human transcriptomic database analysis

To examine how *EDN2* influences the retinal biochemical environment, correlations between *EDN2* and complement components and inflammatory markers (*C1R*, *C1S*, *C1QA*, *C1QB*, *C1QC*, *C2*, *C3*, *C4A*, *C4B*, *CD68*, *GFAP*, *TGFB1*, and *IL-6*) were analyzed using data from the human transcriptomic database GSE115828, focusing on subjects with mild AMD (Mgs levels 1-2; n = 322 samples). The relationship between age and *EDN2* was assessed in these samples. The Pearson correlation coefficients and p values were plotted with ggplot2 (v4.0.1). Relative expressions of *EDN2*, complement elements, *GFAP*, *CD68*, *TGFB1*, and *IL-6* across age groups were plotted with pheatmap (v1.0.13).

### ELISA for EDN2 quantification

Retinal EDN2 levels were quantified using an ELISA kit (E1006Ra, Bioassay Technology Laboratory). Retinal tissue was homogenized, and total protein concentration was determined using a standard assay. EDN2 concentration was measured according to the manufacturer’s protocol, and results were normalized to total protein content and expressed as weight percentage (EDN2 weight relative to total extracted protein weight)

### Intravitreal Injection of AAV or Endothelin Antagonists

To investigate the role of EDN2 in vivo, adeno-associated virus (AAV)-mediated *Edn2* KD was performed in the eyes of 8-month-old SHR and WKY. AAV2 vectors *Edn2*-specific sgRNA (WZ Biosciences Inc.) or control sgRNA were diluted in PBS to 5 x 10^10^ vg/mL. After baseline measurements of IOP, retinal function (ERG), OCT imaging, SBP, and DBP, 1 µL of AAV was injected intravitreally into one eye under anesthesia. Post-treatment assessments were conducted 26-30 days later, followed by tissue collection for transcriptomics, ELISA, and immunohistochemistry (Figure 5A).

Local blockade of EDNRA and EDNRB receptors was achieved through intravitreal injections of BQ123 (ETAR antagonist; Cat. No. B150, Sigma-Aldrich) or BQ788 (ETBR antagonist; 10 μM; Cat. No. B157, Sigma-Aldrich) in SHR and WKY. Baseline IOP and ERG were recorded before the treatment. On Day 0, 1 μL of antagonist or PBS vehicle was injected unilaterally. A second injection was administered on Day 10. IOP was measured on Day 8 and retinal function on Day 9 (post-first injection) and Days 18-19 (post-second injection). Animals were euthanized on Day 20, and injected retinas were harvested for qPCR analysis (Figure 7A). All injections were performed under stereotactic guidance to ensure accurate vitreous delivery, and animals were monitored for signs of ocular inflammation or elevated IOP after treatment.

### Statistical analysis

Data are presented as the mean ± SD. Longitudinal measurements of SBP, DBP, IOP, ERG amplitudes, retinal thicknesses, and retinal arteries and veins lumen areas in SHR and WKY, as well as RGC counts, microglial cell counts, and microglial soma size in 8- and 12-month-old animals, were analyzed by two-way ANOVA with Bonferroni’s multiple comparisons test. Pre- and post-treatment measurements of SBP, DBP, IOP, ERG amplitudes, total retinal thickness, and vessel lumen areas in AAV-injected SHR and WKY were similarly analyzed. qPCR results, αRGC light sensitivity thresholds, and post-treatment RGC/microglial quantification were compared using a two-tailed Student’s t-test. All analyses were performed using Prism 10.0 (GraphPad Software).

### Study approval

All animal experiments were conducted in accordance with the guidelines of the Hong Kong Polytechnic University (PolyU) and approved by the Animal Subjects Ethics Sub-Committee (23-24-893-R-SO-Student).

## Data availability

The raw RNA-sequencing data were uploaded to the GEO repository with accession GSE311105. The values for all data points shown in figures and supplementary figures are available in the Supporting Data Values Excel file.

## Author contributions

Yingkun Cui performed most of the experiments and drafted the manuscript. Man-Ting Au provided guidance on manuscript preparation. Ting Zhang and Feng Pan generated data on light-evoked spike activity of αRGCs. Lanlan Zhang analyzed some of the RNA-sequencing data. Li Pan, Hoi-Lam Li, Margaret MH Wu, Huan Li, Yuqi Zhang, and Shing-Yan Roy Chung conducted part of the in vivo animal studies. Chi-Wai Do and Chunyi Wen initiated and supervised the study, provided resources, secured funding, and wrote and finalized the manuscript.

## Acknowledgments

The authors thank the University Research Facility in Behavioral and Systems Neuroscience at PolyU for their technical support.

## Supplementary Figures

**Figure S1.**
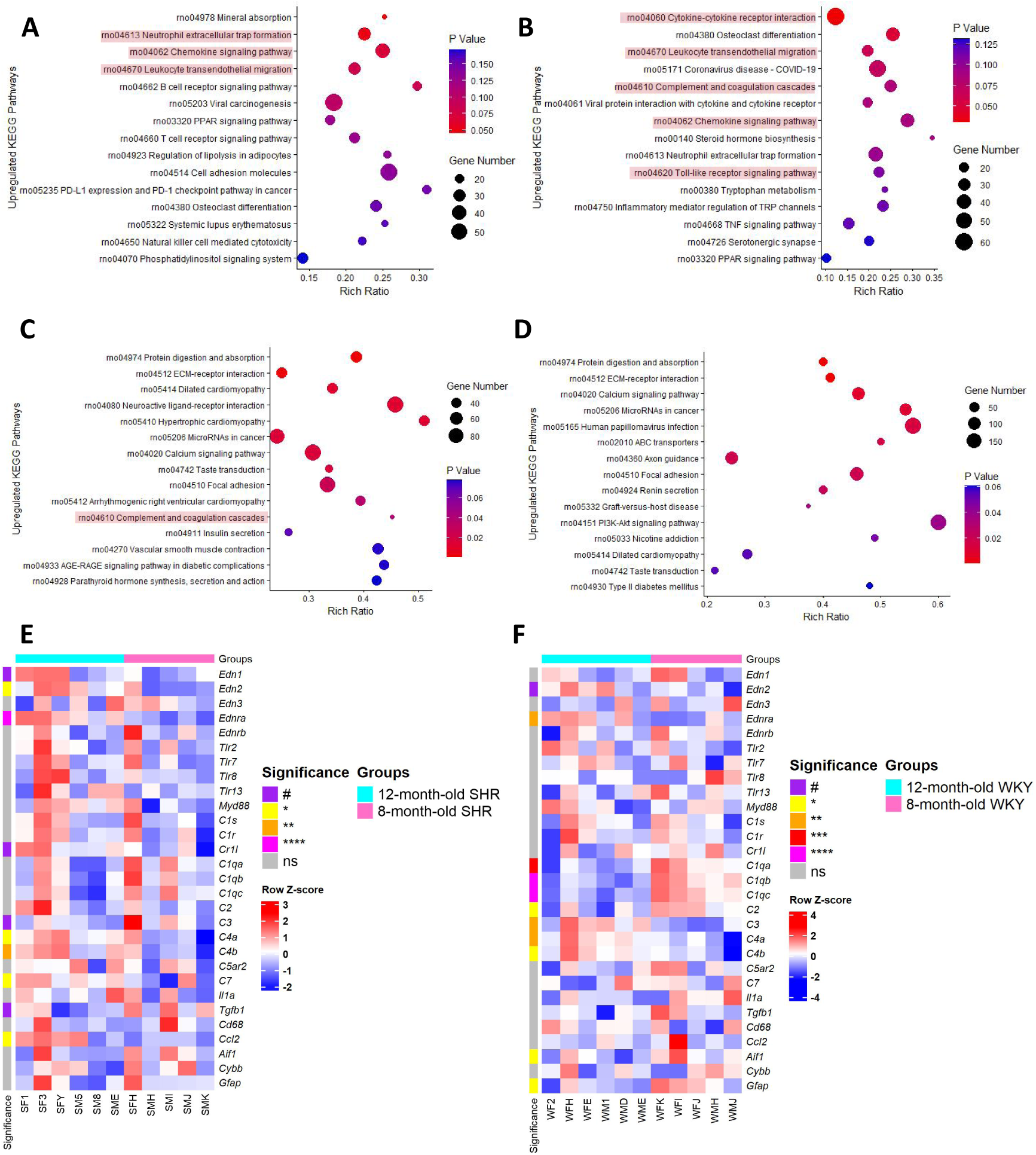
KEGG pathway upregulated by systemic hypertension and aging. **A.** Upregulated KEGG pathways in 8-month-old SHR compared to WKY (n=5 for SHR, n=5 for WKY)**; B.** Upregulated KEGG pathways in 12-month-old SHR versus WKY (n=6 for SHR, n=6 for WKY); **C.** Upregulated KEGG pathways in 12-month-old SHR compared to 8-month-old SHR (n=6 for 12-month-old SHR, n=5 for 8-month-old SHR); **D.** Upregulated KEGG pathways in 12-month-old WKY compared to 8-month-old WKY (n=6 for 12-month-old WKY, n=5 for 8-month-old WKY); **E.** Heatmap of selected target genes comparing 12-month-old SHR against 8-month-old SHR; **F.** Heatmap of selected target genes comparing 12-month-old WKY against 8-month-old WKY.

**Figure S2.**
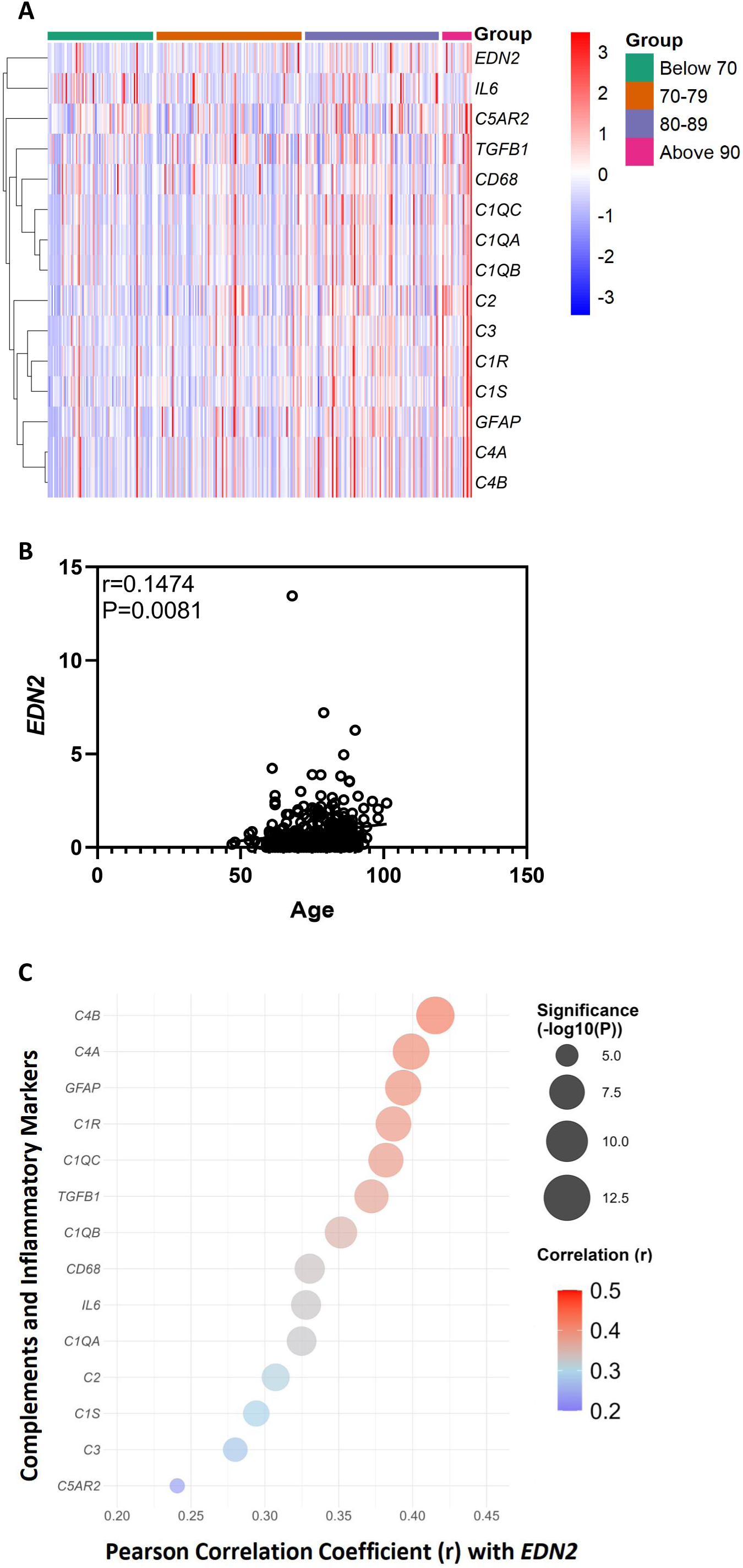
Correlations of *EDN2* with complement factors, inflammatory markers, and aging in human retinas. **A.** Heatmap of EDN2, complement elements, GFAP, CD68, TGFB1, and IL-6 across age groups in 322 human retinas with mild AM; **B.** Correlations between age and EDN2; **C.** Correlations between EDN2 and complement elements, GFAP, CD68, TGFB1, and IL-6 in 322 human retinas with mild AMD.

**Figure S3.**
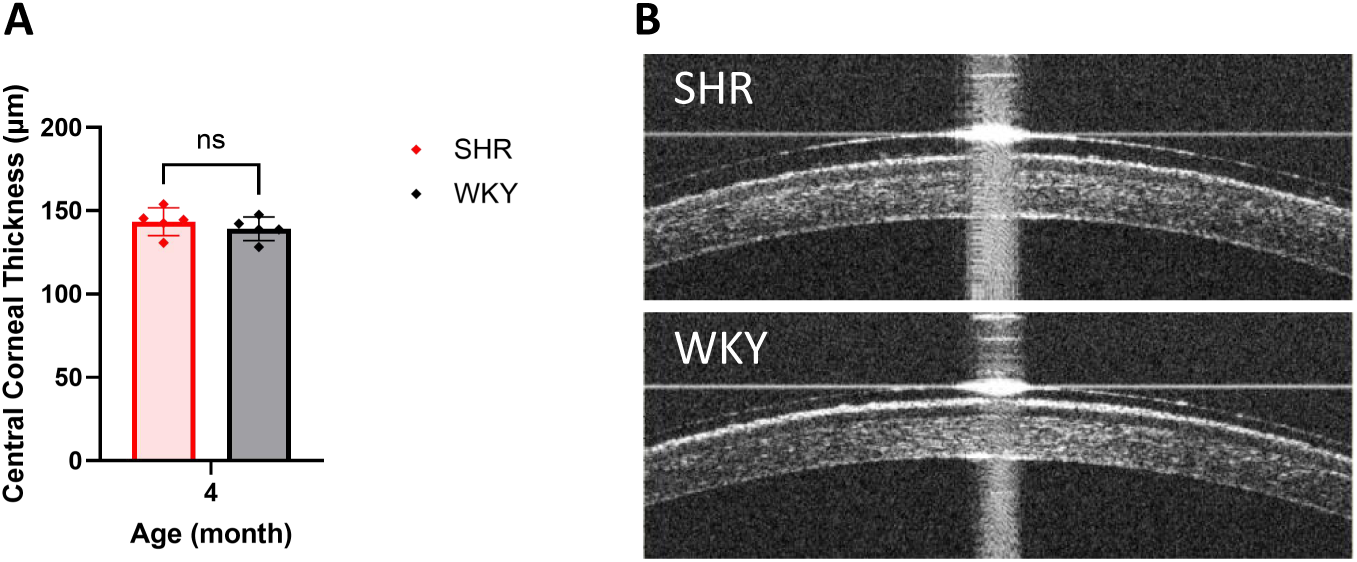
**A.** Measurement of central corneal thickness in 4-month-old SHR and WKY (n=5 for SHR, n=5 for WKY); **B.** Representative images of the central cornea in 4-month-old SHR and WKY. Data are presented as Mean ± SD.

**Figure S4.**
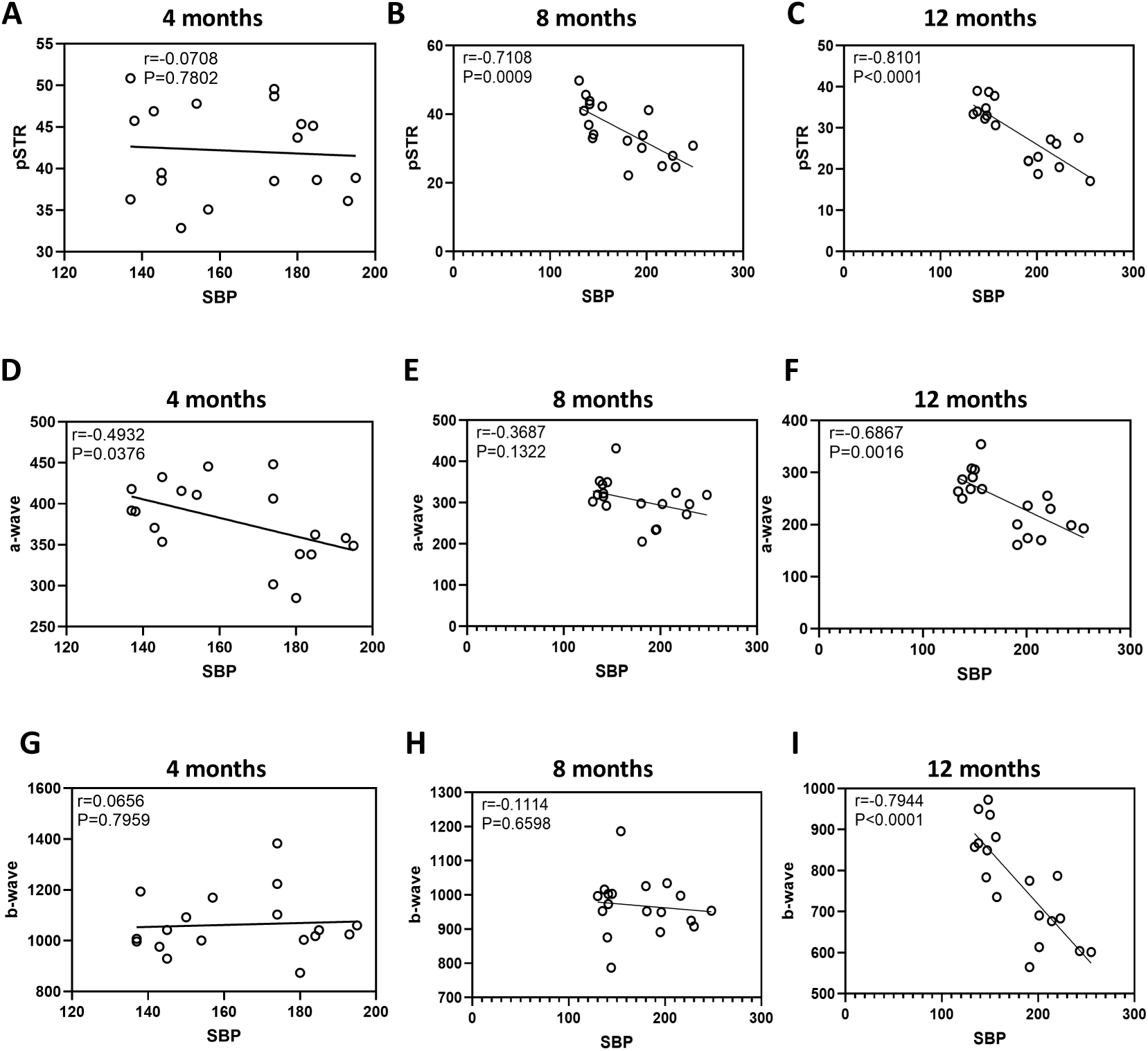
The correlations between SBP and ERG amplitudes at 4, 8, and 12 months. **A-C.** The correlations between SBP and pSTR at 4, 8, and 12 months (n=18); **D-F.** The correlations between SBP and a-wave at 4, 8, and 12 months; **G-I.** The correlations between SBP and b-wave at 4, 8, and 12 months.

**Figure S5.**
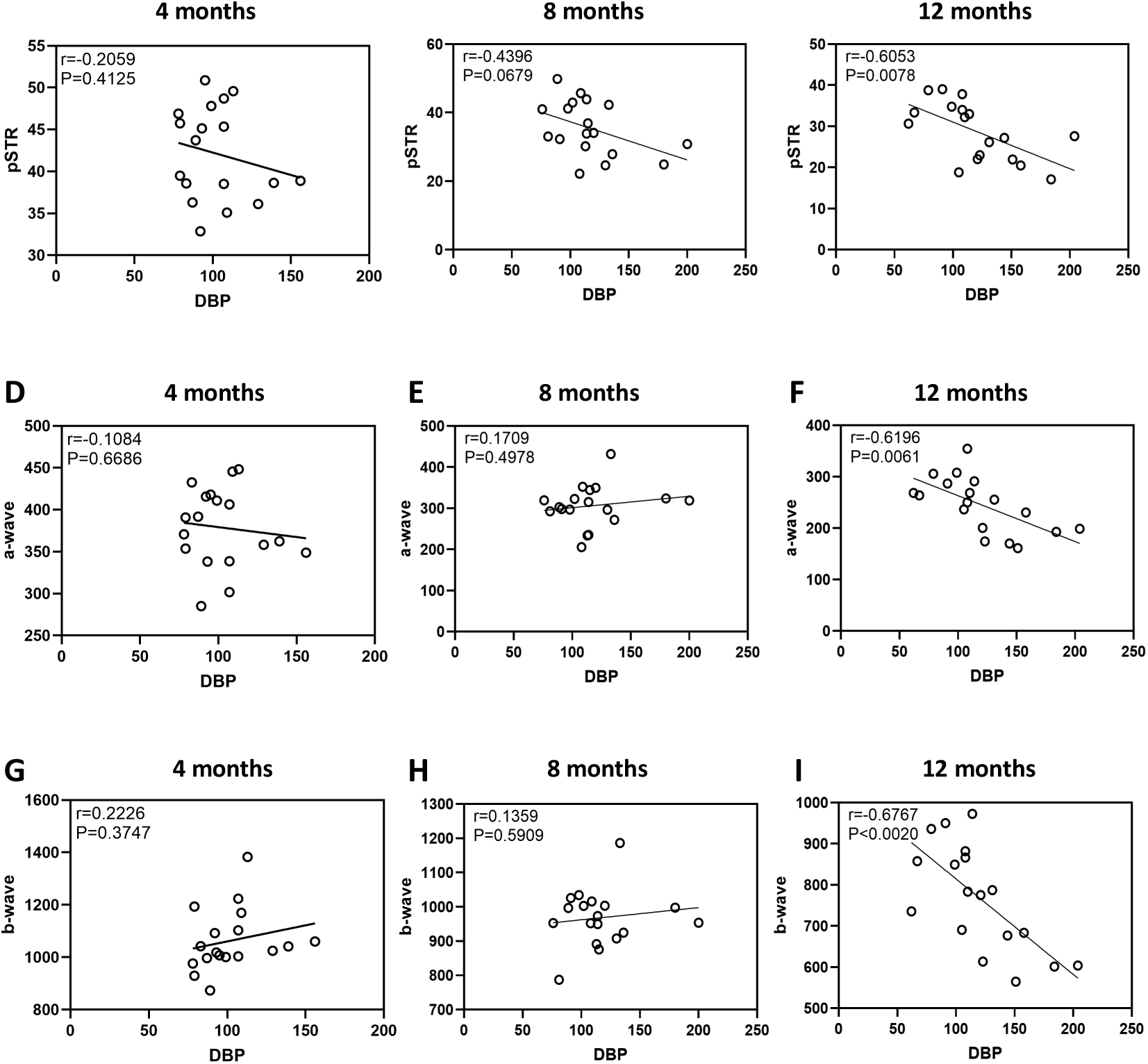
The correlations between DBP and ERG amplitudes at 4, 8, and 12 months. **A-C.** The correlations between DBP and pSTR at 4, 8, and 12 months (n=18); **D-F.** The correlations between DBP and a-wave at 4, 8, and 12 months; **G-I.** The correlations between DBP and b-wave at 4, 8, and 12 months.

